# Repeated morphine exposure activates synaptogenesis and other neuroplasticity-related gene networks in the prefrontal cortex of male and female rats

**DOI:** 10.1101/2020.02.26.966416

**Authors:** Shirelle X. Liu, Mari S. Gades, Andrew C. Harris, Phu V. Tran, Jonathan C. Gewirtz

## Abstract

**Background:** Opioid abuse is a chronic disorder likely involving stable neuroplastic modifications. While a number of molecules contributing to these changes have been identified, the broader spectrum of genes and gene networks that are affected by repeated opioid administration remain understudied.

**Methods:** We employed Next-Generation RNA-sequencing (RNA-seq) to investigate changes in gene expression in adult male and female rats’ prefrontal cortex (PFC) following daily injection of morphine (5.0 mg/kg) for 10 days. Ingenuity Pathway Analysis (IPA) was used to analyze affected molecular pathways, gene networks, and associated regulatory factors.

**Results:** 90% of differentially expressed genes (DEGs) were upregulated in both males and females, with a 35% overlap between sexes. A substantial number of DEGs play roles in synaptic signaling and neuroplasticity. Although broadly similar, some differences were revealed in the gene ontology networks enriched in females and males (e.g., the endocannabinoid pathway in females and neuroinflammation in males).

**Conclusions:** Our results cohere with findings from previous studies based on *a priori* gene selection, while identifying broader gene networks activated by repeated opioid exposure. Our results also reveal novel genes and molecular pathways that are upregulated by repeated morphine exposure.

## 1. Introduction

Opioid addiction is a chronic, often lifelong, disorder, characterized by high rates of relapse (Camí and Farré, 2003). The disorder’s enduring nature is attributable to a variety of long-term changes in neuronal structure and neuronal plasticity (Hyman et al., 2006). These, in turn, are likely induced, and subsequently maintained, through alterations in patterns of gene expression (Nestler, 2001). As such, characterizing the molecular mechanisms underlying opioid addiction is essential for developing more effective treatments.

Investigations into the cellular, molecular, and genetic effects of opioid administration have undergone rapid progress (Imperio et al., 2016; Valentino et al., 2020). Nevertheless, while a number of molecular targets that play key roles in opioid addiction have been identified (Browne et al., 2020; Hurd and O’Brien, 2018), the ways in which a broader range of functional pathways and networks interact to produce and/or maintain addictive behavior remain to be determined. Elucidating molecular networks and targets requires assessing global changes in gene expression after opioid exposure. In an earlier effort along these lines, Spijker et al. (2004) assessed transcriptional changes of 159 genes within pre-selected gene networks, including neurotransmitters, neuronal morphology and plasticity, and intracellular signaling. The advent of Next-Generation RNA Sequencing (RNA-seq) technique allows for measurement of expression levels of genes throughout the transcriptome, without *a priori* selection criteria (Hitzemann et al., 2013). This approach has already yielded critical insights into genomic and downstream regulatory mechanisms underlying cocaine addiction (Walker et al., 2018). The current study is the first to apply RNA-seq to investigate the transcriptional changes following repeated opioid (morphine) exposure.

The role of dysregulated brain reward pathways in addiction has long been recognized (Kreek and Koob, 1998). Both clinical and preclinical studies have implicated alterations in prefrontal cortex (PFC) function as critically important in producing compulsive drug use, drug seeking, and relapse (Goldstein and Volkow, 2002; Kalivas et al., 2005; Koya et al., 2009; Wilson et al., 2004). Responding to reward or in anticipation of reward in a variety of contexts is a function of the activity of glutamatergic projections from PFC subregions to the nucleus accumbens (NAc). Activity in both structures is modulated by dopaminergic inputs from the ventral tegmental area (Hyman et al., 2006). The PFC is therefore an appropriate locus for investigating changes in gene expression induced by exposure to opioid drugs.

In the current study, we tested global gene expression changes in the PFC 24 hours after repeated, daily injection of morphine This is a time point after opioid exposure at which rats are prone to exhibit anhedonia and relapse to drug-seeking (Rothwell et al., 2009; Shaham et al., 1996). Furthermore, we recently reported that the intensity of morphine withdrawal-induced anhedonia predicts the severity of subsequently acquired morphine self-administration (Swain et al., 2019). Hence, our approach has the potential to identify genes and gene ontology networks that are important in vulnerability to opioid addiction. In view of differences in vulnerability of males and females to this disorder (Becker et al., 2017; Brady and Randall, 1999; Lee and Ho, 2013), and the historical paucity of studies in this area that have included female subjects or been powered sufficiently to measure sex differences (Becker and Koob, 2016), both male and female groups were included. Commonalities and differences between the sexes in the effects of repeated opioid exposure on genomic regulation could therefore be characterized.

## 2. Materials and Methods

### 2.1 Animals

Thirty-six adult Sprague Dawley rats (18 males, 18 females, 65-75 days) were obtained from Envigo (Indianapolis, IN). Animals were same-sex pair-housed in polypropylene cages with *ad libitum* access to food and water. The colony was maintained on a 12-h light-dark cycle. All procedures were conducted during the light phase of the light-dark cycle. Animals were habituated for at least 1 week before experimentation. All animal protocols were approved by the Institutional Animal Care and Use Committee (IACUC) of the University of Minnesota.

### 2.2 Drug treatments

Animals were randomly assigned to morphine or saline conditions, which resulted in 9 animals/group (Male/Morphine, Male/Saline, Female/Morphine, and Female/Saline). Morphine sulfate (Research Triangle International, NC) was dissolved in 0.9% sterile saline and injected subcutaneously. Animals received daily injections of either morphine (5.0 mg/kg) or an equivalent volume of saline daily over 10 consecutive days. This is similar to the dosing regimen used in previous studies of behavioral sensitization to the effects of morphine (Paolone et al., 2003; Rothwell et al., 2010; Shippenberg et al., 1996).

### 2.3 Tissue collection

Animals were sacrificed 24 hours after the last injection. Prefrontal cortex was dissected bilaterally, immediately flash-frozen in liquid nitrogen, and stored at −80□. Samples within each condition for each sex were randomly pooled in groups of three prior to RNA preparation.

### 2.4 RNA preparation and sequencing

RNA-seq was conducted as previously described (Barks et al., 2018). Total RNA was isolated using the RNeasy Mini Kit (Qiagen) according to the manufacturer’s protocol. Isolated RNA was further purified and concentrated using the MinElute Cleanup Kit (Qiagen). Library preparation and RNA-seq were conducted at the University of Minnesota Genomics Center. RNA was quantified using the RiboGreen RNA Assay kit (Invitrogen) and assessed for quality using capillary electrophoresis (Agilent BioAnalyzer 2100; Agilent). Barcoded libraries were constructed for each sample using the TruSeq RNA v2 kit (Illumina). Libraries were size-(200 bp) selected and sequenced (50bp paired-end read, ~20 million reads/library) using Illumina HiSeq 2500.

### 2.5 RNA-seq analysis

Quality control on raw sequence data was performed with FastQC. Mapping of reads was performed via Hisat2 (version 2.1.0) using the rat genome (rn6) as reference. Differentially expressed genes (DEGs) were identified by gene-wise negative binomial generalized linear models using the EdgeR feature in CLC Genomics Workbench (Qiagen, version 10.1.1). The generated list was filtered based on ≥ 2x absolute fold change and false discovery rate (FDR) corrected *p*-value (*q*-value) < 0.05. Principle Component Analysis (PCA) of DEGs was conducted via unsupervised clustering.

### 2.6 Ingenuity Pathway Analysis

DEGs were annotated by Ingenuity pathway Analysis (IPA; Qiagen) to identify relevant canonical pathways, molecular networks and cellular functions that showed significant alterations in experimental versus control groups as previously described (Barks et al., 2018). Statistical significance [*p* < 0.05; −log(*p*) > 1.3] was determined by Fisher’s exact test.

### 2.7 Real-time quantitative PCR (RT-qPCR)

RT-qPCR was performed on technical replicates (6 samples/group) as previously described (Barks et al., 2018). RNA was isolated using RNAqueous Total RNA Isolation Kit (Invitrogen). cDNA synthesis was performed using High-Capacity RNA-to-cDNA Kit (Applied Biosystems). qPCR was performed using a TaqMan Universal PCR Master Mix (Applied Biosystem) and TaqMan gene expression assays (ThermoFisher Scientific) on a DNA analyzer (QuantStudio 3, ThermoFisher Scientific). Beta actin (*Actb*) was used as an endogenous control. Samples were run in duplicate, normalized to *Actb* and averaged to generate fold-changes relative to controls. Results were analyzed by two-tailed *t*-tests in RStudio (version 1.2.5033), with α set at *p* < 0.05.

## 3. Results

### 3.1. Repeated morphine injections altered gene expression in the PFC

NGS data were aligned to 17,336 loci in morphine- or saline-treated rats (Figs. 1A, 1B). Principal Component Analysis of the 500 most divergent genes showed that samples clustered into four discrete groups based on drug treatment and sex (Supplementary Material, S.1). In male rats, 377 genes were differentially expressed in the morphine-treated relative to the saline-treated group, among which 337 (89%) were up-regulated and 40 (11%) were down-regulated (Supplementary Material, S.2). In female rats, 409 genes were significantly differentially expressed in morphine-treated relative to saline-treated rats, with 370 (90%) up-regulated and 39 (10%) down-regulated (Supplementary Material, S.3). RNA-seq results were verified by RT-qPCR with selected genes that are known to regulate nervous system development and function (Figs. 1C, 1D; Supplementary Material, S.4).

**Fig. 1.**
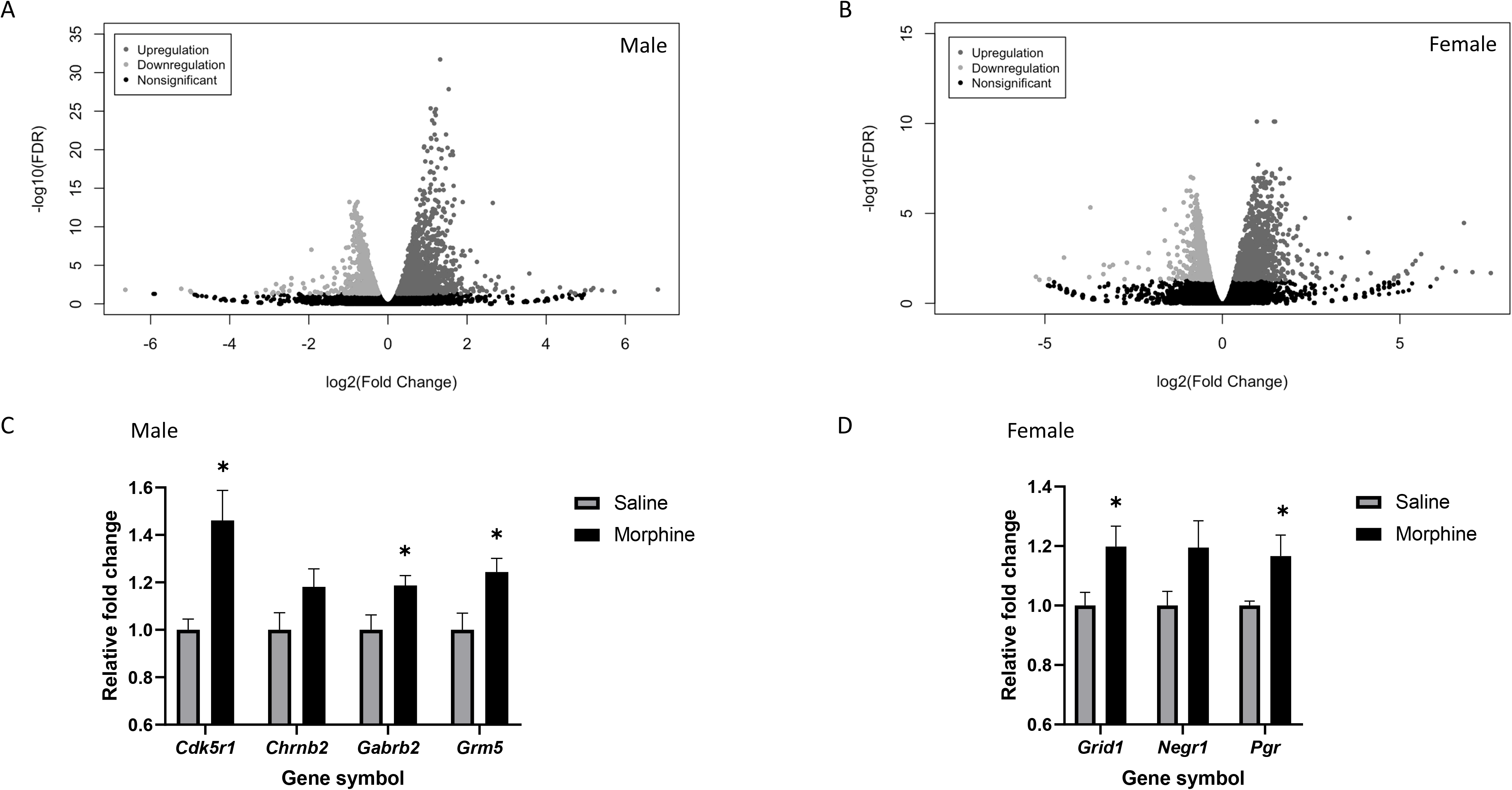
RNA-seq and RT-qPCR validation of transcriptional changes in the rat PFC following a 10-day morphine injection regimen. (A, B) Volcano plots generated from unfiltered RNA-seq data from males and females, respectively [FDR, False Discovery Rate corrected *p*-value (*q*-value)]. (C, D) RT-qPCR validation of male and female RNA-seq datasets (*n* = 5-6/group. * *p* < 0.05 compared with the respective saline control group).

### 3.2 Differentially expressed genes implicated changes in neuronal plasticity and intra-/inter-cellular signaling pathways induced by repeated morphine exposures

Male and female groups shared a subset of 204 (35%) DEGs (Fig. 2A). This overlapping set of genes was analyzed using IPA to identify non-sex-specific effects. DEGs were enriched in canonical pathways critical for synaptic/intracellular signaling, including synaptogenesis, longterm potentiation, opioid signaling, dopamine-DARPP32 signaling and ephrin receptor signaling pathways (Fig. 2B). All of the top affected pathways were activated (z-score ≥ 2).

**Fig. 2.**
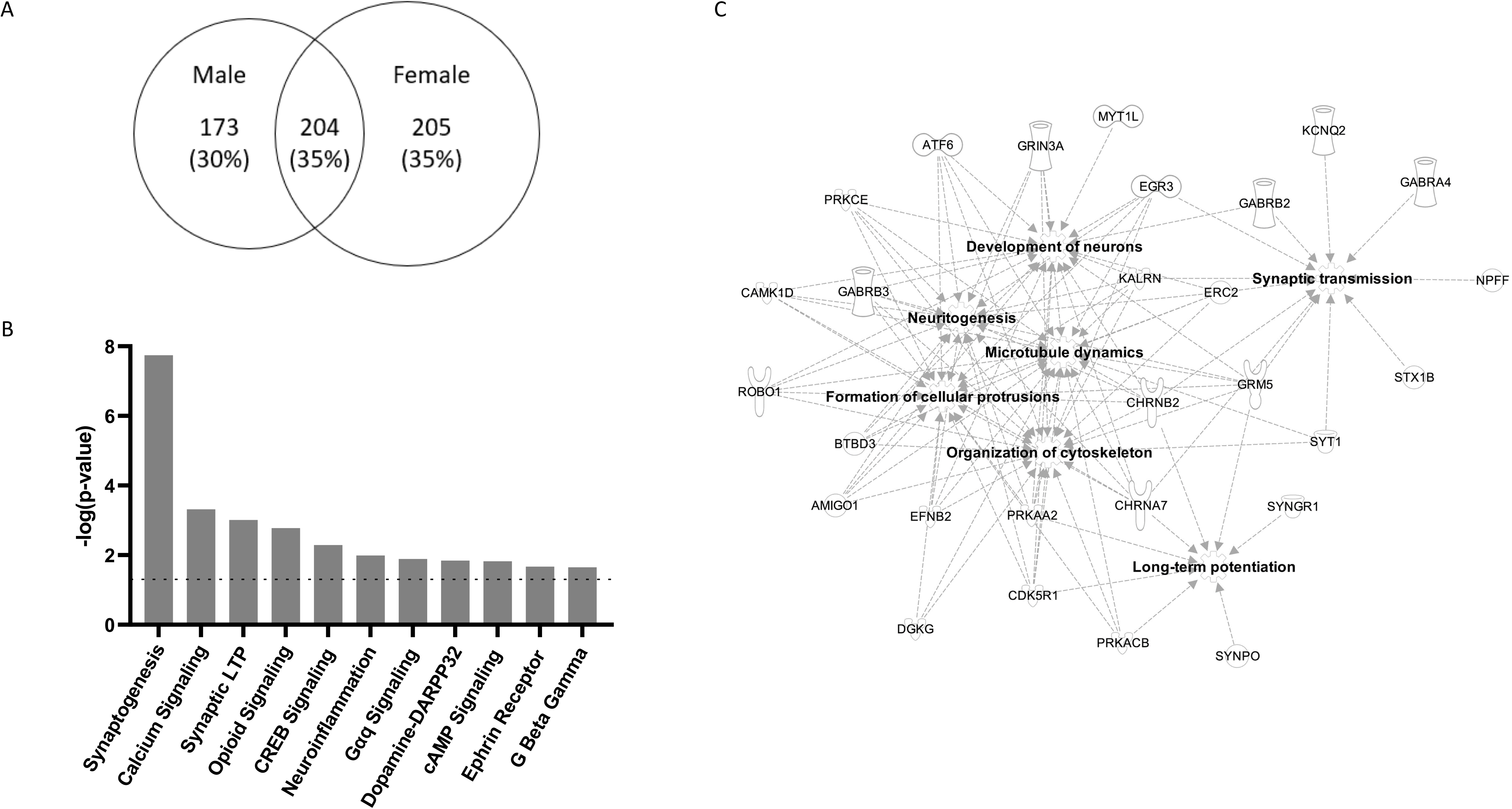
IPA analysis of overlapping genes, affected canonical pathways and gene networks. (A) Venn diagram of filtered total number of DEGs in males and females. (B) DEGs in both males and females were enriched in cell signaling-, and synaptic growth- and plasticity-related pathways [absolute *z*-score ≥ 2; *p*-value < 0.05, −log(*p*-value) > 1.3, dotted line]. (C) Annotated functions of gene networks in PFC affected in all morphine-treated rats (absolute *z*-score ≥ 2; *p*-value < 0.05).

DEGs that were shared between sexes were also analyzed via IPA to identify enrichment of diseases and functions (Fig. 2C). Enriched functional groups of genes included those coding for neurotransmitter receptor subunits (*Chrna7, Chrnb2, Gabra4, Gabrb2, Gabrb3, Grin3a, Grm5, Htr2a, Htr5a, Pgr*), intra-/inter-cellular signaling regulation (*Camk2d, Cdk5r1, Efnb2, Kalrn, Prkaa2, Prkacb, Prkce*), and synaptic morphology/function (*Stx1b, Syngr1, Synpo, Syt1*) (Supplementary Materials, S.2 and S.3). In addition, these DEGs implicated JAK1/2, FEV, and ADORA2A as key upstream regulators of the transcriptional effects of repeated morphine exposure (Supplementary Material, S.5).

Finally, a comparison of separate analyses of DEGs in each sex showed enrichment of the neuroinflammatory pathway in males only and the endocannabinoid pathway and long-term depression pathway in females only (Supplementary Materials, S.6 and S.7).

## 4. Discussion

We analyzed the transcriptome from the rat PFC, a key brain structure implicated in addictive behavior, to identify *a posteriori* genes and gene networks dysregulated by repeated opioid exposure. The most striking outcome at a global level was that 90% of DEGs were upregulated in both sexes. Such a high degree of upregulation is consistent with the results of a previous RT-qPCR study conducted on the NAc (Spijker et al., 2004) at the same time point after drug exposure and implicates widespread transcriptional activation as a regulatory response following chronic opioid exposure.

The overlapping upregulated genes common to both sexes participate in canonical pathways broadly associated with synaptogenesis and neuroplasticity, consistent with the idea that recruitment of these cellular processes is a fundamental process in the development of addiction (Lüscher and Malenka, 2011). Altered PFC gene expression indicates activation in molecular networks associated with opioid and dopamine signaling, and intracellular Ca^2+^, G-protein, and cAMP/CREB signaling pathways. These findings support the view that addiction is induced via neuroadaptations and neuroplasticity and concomitant alterations in opioidergic and dopaminergic neurotransmission via G-protein-coupled receptors (Dani et al., 2001; Kauer and Malenka, 2007). For example, the cAMP-dependent pathway is upregulated by chronic morphine treatment (Avidor-Reiss et al., 1996; Nestler and Tallman, 1988; Terwilliger et al., 1991), resulting in the activating phosphorylation of CREB (Haghparast et al., 2014; Morón et al., 2010). These changes in turn likely affect neurotransmitter release and enhance synaptic connectivity (Chavez-Noriega and Stevens, 1994; Weisskopf et al., 1994). It has been further proposed that such upregulation of the cAMP pathway represents a compensatory response to the acute inhibitory effect of opioid administration (Sharma et al., 1977, 1975; Traber et al., 1975), that may play an important role in opioid dependence, tolerance, and withdrawal (Hamdy et al., 2001; Lai et al., 2014; Nestler, 2016).

A number of the genes that were significantly upregulated in both sexes have already been implicated in addiction, validating this study’s approach. These include ionotropic and metabotropic glutamate receptors (*Grin3a, Grm5*), nicotinic (*Chrna7, Chrnb2*) and serotonergic (*Htr2a, Htr5a*) receptor subtypes, and the progesterone receptor (Evans and Foltin, 2006; Jackson et al., 2006; Muneoka et al., 2010; Popik and Wróbel, 2002; Yuan et al., 2013). In addition, our findings reveal potential novel molecular substrates of opioid-induced addiction-related plasticity. These include cyclin dependent kinase 5 (Cdk5) regulatory subunit 1 (*Cdk5r1*) and ephrin B2 *(Efnb2). Ck5r1* encodes a neuronal-specific activator of Cdk5, which is strongly implicated in addictive properties of cocaine and opioids (Bibb et al., 2001; Ferrer-Alcón et al., 2003; Narita et al., 2005). Cdk5r1 upregulation in this study differs from a previous study that found downregulation of Cdk5/Cdk5r1 in morphine-treated rats (Ferrer-Alcón et al., 2003). These disparate findings may reflect differences in the timing of the assay relative to drug exposure (Spijker et al., 2004).

The Ephrin family of tyrosine kinase-related receptors regulates neurogenesis, neuronal migration, synaptic plasticity, axon guidance and neuroadaptation (Ashton et al., 2012; McClelland et al., 2009; Xiao et al., 2006). Ephrins also interact with glutamatergic and dopaminergic pathways (Essmann et al., 2008; Piccinin et al., 2010; Planagumà et al., 2016; Yue et al., 1999). Despite participating in such a highly relevant set of functional domains, the possible role of ephrins in the pathophysiology of mental illnesses, including addiction, has received relatively little attention. Nevertheless, the current finding of increased *Efnb2* expression in the PFC adds to two previous transcriptomic screens that detected its upregulation in the NAc after repeated morphine exposure (Martínez-Rivera et al., 2019; Spijker et al., 2004). Thus, accumulating evidence implicates *Efnb2* in the long-term effects of opioid exposure in the meso-corticolimbic system.

Analysis of upstream regulators from the common DEGs of both sexes suggests increased activity of JAK1/2, FEV, and ADORA2A. Both FEV and ADORA2A are potential hubs in the network of genes involved in opioid addiction. FEV transcription factor (PET-1) is localized in serotonin neurons in the CNS, where it is necessary for serotonin synthesis and release (Liu et al., 2010; Puzerey et al., 2015; Wyler et al., 2016). Our data additionally showed that genes coding the serotonin 5-HT2a and 5-HT5a receptors were upregulated after opioid exposure further supporting the possibility of interactions between the opioidergic and serotonergic systems in the PFC (Marek and Aghajanian, 1998; Marek et al., 2001). The adenosine 2a receptor (ADORA2A) also interacts with the opioid system (Brown et al., 2009; Yao et al., 2006) and forms heteromeric complexes with other receptors, including the dopaminergic D2 receptor and the glutamatergic mGluR5 receptor (Ferré et al., 2007), which was upregulated in our study. Hence, the emergence of PET-1 and the adenosine 2a receptor as upstream regulators may provide insights into the hubs underlying gene networks critical in the development or maintenance of opioid addiction.

Our identification of molecular networks that were differentially affected by morphine in males and females highlights potential mechanisms underlying the gender-specific effects of vulnerability to opioid addiction. Female rats showed significant enrichment of genes in the long-term depression and endocannabinoid synaptic pathways. The latter likely reflects the interaction between cannabinoid and opioid systems (Ledent et al., 1999; Martin et al., 2000; Wenzel and Cheer, 2018), which occurs in response to opioid drugs (Maldonado and Rodríguez De Fonseca, 2002). Additionally, endocannabinoid receptors participate in estradiol-potentiated cocaine-induced locomotor activity (Peterson et al., 2016). The interactions among endocannabinoid, opioid and hormonal systems may represent female-specific mechanisms underlying addiction vulnerability (Anker and Carroll, 2011; Carroll and Anker, 2009; Chartoff and McHugh, 2016; Lynch et al., 2002). In contrast, male rats showed enrichment of genes in neuroinflammatory pathways, adding to an emerging literature on the role of neuroinflammatory processes in sex differences in the effects of opioids (Averitt et al., 2019; Doyle et al., 2017).

In summary, repeated morphine administration produced substantial upregulation of gene transcription in the PFC, particularly affecting genes involved in synaptogenesis, neuroplasticity, and neuronal development. These changes may reflect a compensatory response to repeated stimulation of mu opioid receptors. It is noteworthy that assessment of transcriptional changes 24-hr after drug exposure in this study coincides with robust manifestations of opioid withdrawal symptoms (Bechara et al., 1995; Gold et al., 1994; Hand et al., 1988; Rothwell et al., 2009). We have found that the intensity of one of these measures - withdrawal-induced anhedonia - correlates with the severity of subsequent morphine self-administration. Thus, to the extent that their functional relevance is confirmed, a subset of the genes and gene ontology pathways identified in this study may represent opioid-stimulated biomarkers for long-term vulnerability to opioid addiction.

## Author Disclosures

### Contributors

SXL contributed to analysis and interpretation of RNA-seq data, execution of RT-qPCR, and primary writing and preparation of the manuscript. MSG contributed to study design and collection and analysis of RNA-seq data. ACH contributed to hypothesis generation and study design. PVT contributed to processing of RNA-seq and RT-qPCR samples and analysis of RNA-seq data. JCG contributed to hypothesis generation, study design, interpretation of RNA-seq data, primary writing of manuscript, and project oversight. All authors contributed to manuscript revision.

## Acknowledgments

We thank Amanda Barks for help with RT-qPCR and Dr. Juan E. Abrahante Lloréns for help with data analysis.

