## Supplementary figures and images for "Repeated morphine exposure activates synaptogenesis and other neuroplasticity-related gene networks in the prefrontal cortex of male and female rats"

### Supplementary Material S.1

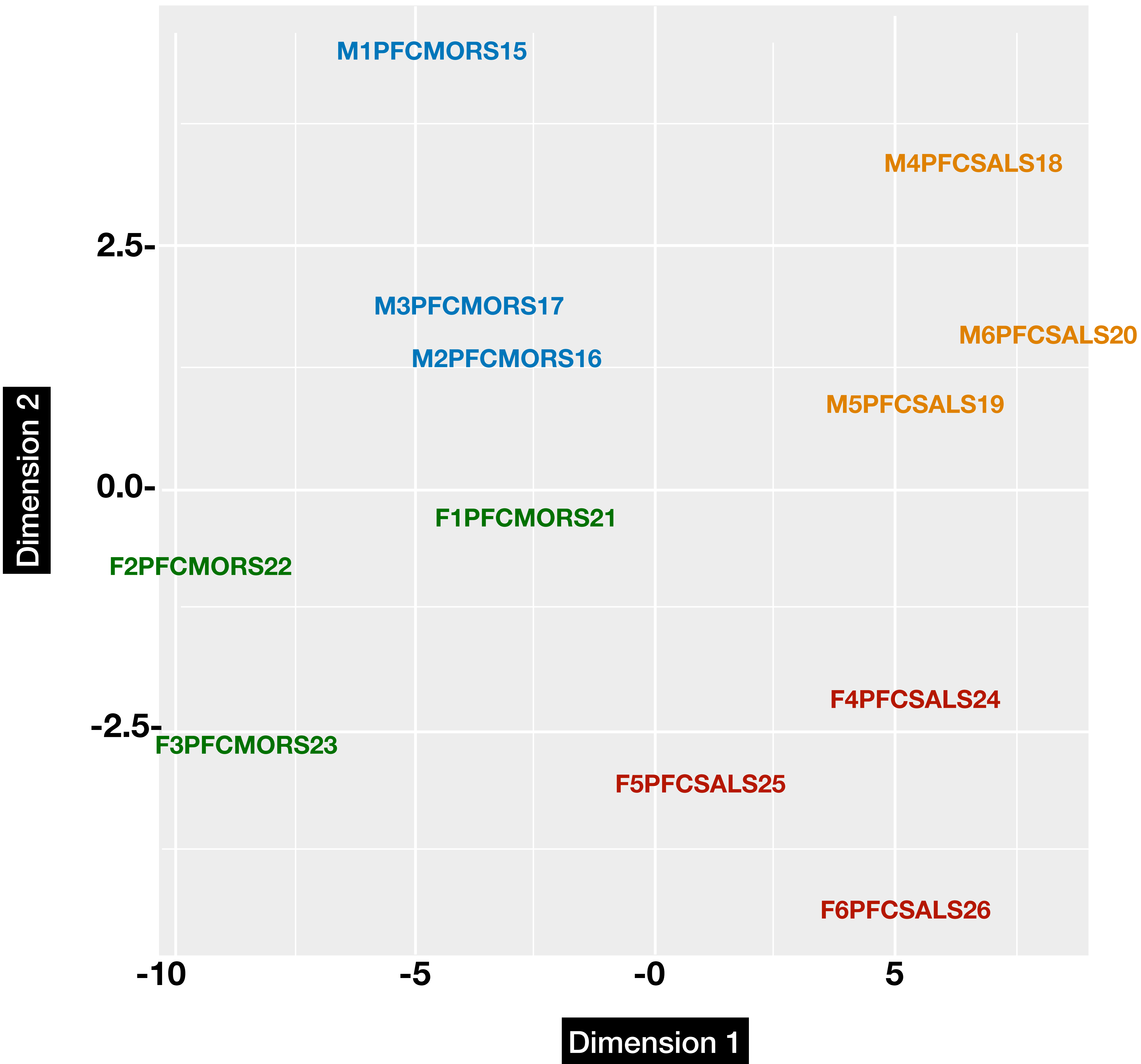
