## Supplementary Material S.2 for "Repeated morphine exposure activates synaptogenesis and other neuroplasticity-related gene networks in the prefrontal cortex of male and female rats"

| Gene Symbol | Fold Change | FDR-adjusted p-value (q-value) |
|---|---|---|
| *Abcb7* | 2.090 | 2.070E-03 |
| *Ackr2* | 2.562 | 2.806E-04 |
| *Actr2* | 2.311 | 3.362E-25 |
| *Acvr1b* | 2.219 | 2.136E-12 |
| *Adam23* | 2.601 | 5.892E-10 |
| *Ager* | -2.042 | 1.719E-03 |
| *Agps* | 2.010 | 3.293E-02 |
| *Alg10* | 2.778 | 5.776E-03 |
| *Amigo1* | 2.488 | 1.420E-05 |
| *Ankle2* | 2.099 | 1.195E-03 |
| *Ankrd34c* | 2.080 | 1.794E-02 |
| *Ankrd52* | 2.247 | 2.459E-02 |
| *Ap2b1* | 2.238 | 2.567E-10 |
| *Arhgap26* | 2.092 | 4.116E-02 |
| *Armc6* | 2.087 | 6.875E-04 |
| *Arrdc3* | 3.144 | 2.566E-07 |
| *Atf2* | 2.018 | 3.602E-11 |
| *Atf6* | 2.063 | 9.056E-04 |
| *Atic* | 2.068 | 7.644E-04 |
| *Atl3* | 2.154 | 1.638E-02 |
| *Atp2b4* | 3.169 | 2.987E-12 |
| *B3galt1* | 2.828 | 6.249E-07 |
| *Bc1* | -2.572 | 2.298E-05 |
| *Bcl2l13* | 3.087 | 4.511E-05 |
| *Brms1* | -7.585 | 4.701E-02 |
| *Btbd3* | 2.261 | 2.391E-10 |
| *C1galt1* | 2.138 | 2.933E-02 |
| *C2cd4c* | 2.278 | 6.934E-05 |
| *Cadps* | 2.133 | 3.975E-22 |
| *Camk1d* | 2.572 | 5.020E-12 |
| *Camk2g* | 2.507 | 3.019E-02 |
| *Casc3* | 2.276 | 7.544E-08 |
| *Ccdc132* | 2.718 | 3.897E-08 |
| *Ccdc173* | -2.064 | 3.852E-02 |
| *Ccser1* | 2.949 | 6.467E-03 |
| *Cd34* | 2.345 | 4.281E-14 |
| *Cdh19* | 3.017 | 1.155E-03 |
| *Cdk5r1* | 2.475 | 1.489E-07 |
| *Cds1* | 2.416 | 6.279E-14 |
| *Cela3b* | -2.896 | 5.169E-03 |
| *Chid1* | 15.083 | 2.515E-02 |
| *Chrm1* | 2.010 | 3.208E-14 |
| *Chrm2* | 3.646 | 4.749E-04 |
| *Chrna7* | 2.148 | 1.191E-02 |
| *Chrnb2* | 2.554 | 7.195E-09 |
| *Chst15* | 2.376 | 4.838E-04 |
| *Cldn3* | -2.049 | 4.288E-02 |
| *Clic2* | -2.272 | 1.092E-02 |

| | | |
|---|---|---|
| *Clint1* | 2.165 | 8.257E-06 |
| *Clvs2* | 2.256 | 1.174E-02 |
| *Cmip* | 2.339 | 4.822E-22 |
| *Coro2a* | 3.139 | 7.411E-06 |
| *Cort* | -2.142 | 2.084E-05 |
| *Cpeb4* | 2.400 | 8.365E-08 |
| *Ctdspl2* | 2.388 | 1.648E-04 |
| *Cul5* | 2.006 | 5.175E-04 |
| *Cyth1* | 2.229 | 9.695E-06 |
| *Dcbld2* | 2.022 | 3.015E-03 |
| *Dck* | 2.057 | 2.896E-04 |
| *Dctn4* | 2.488 | 1.128E-07 |
| *Ddah1* | 2.094 | 1.031E-11 |
| *Dgkg* | 2.351 | 3.481E-07 |
| *Dhcr24* | 2.435 | 1.142E-09 |
| *Dhx33* | 2.255 | 5.131E-04 |
| *Diras2* | 2.348 | 1.706E-12 |
| *Dlc1* | 2.424 | 2.386E-05 |
| *Dlk1* | -2.114 | 1.949E-02 |
| *Dnajc16* | 2.755 | 1.394E-11 |
| *Dnm1l* | 2.091 | 3.604E-15 |
| *Dusp23* | 2.694 | 1.042E-02 |
| *Dusp8* | 2.724 | 1.268E-09 |
| *Ece1* | 2.303 | 7.411E-06 |
| *Edem3* | 2.434 | 1.853E-03 |
| *Ednrb* | 2.512 | 2.637E-04 |
| *Efna5* | 2.090 | 4.127E-02 |
| *Efnb2* | 6.279 | 8.064E-14 |
| *Egr3* | 2.776 | 3.395E-06 |
| *Eif4enif1* | 2.190 | 6.482E-07 |
| *Elovl7* | 2.196 | 5.177E-03 |
| *Ephb2* | 2.061 | 1.394E-03 |
| *Erc2* | 2.701 | 5.033E-08 |
| *Esrrg* | 2.201 | 9.105E-04 |
| *Exoc4* | 2.422 | 1.468E-10 |
| *Exoc8* | 2.160 | 6.066E-04 |
| *Fads6* | 2.003 | 9.421E-08 |
| *Fam126a* | 2.791 | 1.708E-02 |
| *Fam13a* | 2.106 | 1.247E-02 |
| *Fam163b* | 2.072 | 1.547E-08 |
| *Fam168b* | 2.248 | 3.962E-24 |
| *Fam219a* | 2.084 | 8.461E-18 |
| *Fam49a* | 2.313 | 5.895E-06 |
| *Fbxl16* | 2.125 | 3.323E-16 |
| *Fgf14* | 2.269 | 7.027E-04 |
| *Ficd* | 2.581 | 2.384E-04 |
| *Fnbp1l* | 2.054 | 2.578E-04 |
| *Foxo1* | 2.676 | 3.119E-04 |
| *Foxred2* | 3.212 | 5.257E-03 |

| | | |
|---|---|---|
| *Frmpd4* | 2.699 | 1.274E-12 |
| *Fry* | 2.154 | 1.194E-12 |
| *Fto* | 2.199 | 4.633E-13 |
| *Gabra3* | 2.189 | 1.718E-09 |
| *Gabra4* | 2.551 | 1.318E-19 |
| *Gabrb2* | 2.286 | 2.442E-03 |
| *Gabrb3* | 2.426 | 5.537E-10 |
| *Gabrg2* | 2.130 | 4.426E-19 |
| *Galc* | 2.186 | 1.838E-04 |
| *Galntl6* | 2.072 | 9.768E-04 |
| *Gda* | 2.513 | 7.790E-10 |
| *Glra2* | 2.542 | 2.877E-02 |
| *Glyr1* | 2.464 | 1.738E-15 |
| *Gna11* | 2.362 | 1.547E-08 |
| *Gnl3l* | 2.419 | 1.715E-08 |
| *Golga7b* | 2.107 | 3.866E-09 |
| *Golph3l* | 2.003 | 3.563E-06 |
| *Gpd1l* | 2.985 | 5.210E-08 |
| *Gpr165* | 2.938 | 5.087E-03 |
| *Gpr17* | 3.505 | 1.904E-03 |
| *Grid1* | 2.658 | 8.121E-08 |
| *Grin3a* | 2.240 | 2.137E-02 |
| *Grm5* | 2.326 | 4.838E-07 |
| *Gstt2* | 2.279 | 2.417E-02 |
| *Gtf2a1* | 2.061 | 4.431E-02 |
| *Guca2a* | -5.425 | 4.512E-04 |
| *Hif1an* | 2.275 | 1.226E-02 |
| *Hnrnpr* | 2.602 | 2.312E-05 |
| *Hp* | -2.041 | 6.925E-03 |
| *Hrh1* | 3.634 | 6.218E-04 |
| *Hs6st2* | 2.155 | 6.511E-04 |
| *Hsbp1l1* | -2.593 | 2.475E-02 |
| *Hsd17b1* | -2.153 | 3.730E-02 |
| *Hsp90aa1* | 2.922 | 2.199E-04 |
| *Hspa13* | 2.042 | 5.362E-08 |
| *Hspa9* | 2.112 | 4.296E-26 |
| *Htr1a* | 3.467 | 2.332E-03 |
| *Htr2a* | 3.629 | 4.758E-07 |
| *Htr5a* | 2.200 | 9.095E-04 |
| *Hunk* | 2.654 | 8.035E-03 |
| *Il10ra* | 2.580 | 1.971E-02 |
| *Il1r1* | 2.741 | 1.753E-03 |
| *Inpp5b* | 2.372 | 1.804E-02 |
| *Ipcef1* | 3.296 | 2.503E-06 |
| *Irs1* | 2.359 | 6.937E-03 |
| *Itgb8* | 3.008 | 2.473E-03 |
| *Jph1* | 2.439 | 1.315E-04 |
| *Kalrn* | 2.124 | 3.031E-22 |
| *Kbtbd4* | 2.069 | 2.967E-05 |

| | | |
|---|---|---|
| *Kcna1* | 2.459 | 6.542E-03 |
| *Kcnf1* | 2.161 | 2.218E-10 |
| *Kcnj2* | 4.620 | 2.176E-03 |
| *Kcnj3* | 2.111 | 2.335E-09 |
| *Kcnk9* | 2.733 | 1.370E-02 |
| *Kcnq2* | 2.117 | 1.430E-04 |
| *Kcns2* | 2.378 | 1.500E-03 |
| *Kctd21* | 2.213 | 1.939E-03 |
| *Kit* | 2.128 | 1.393E-06 |
| *Klhl12* | 2.242 | 1.718E-09 |
| *Kpna1* | 2.108 | 1.026E-05 |
| *Kpna3* | 2.004 | 2.131E-09 |
| *Kpnb1* | 2.224 | 9.707E-18 |
| *Kremen1* | 2.465 | 1.402E-02 |
| *Lhfpl4* | 2.689 | 1.158E-03 |
| *Lims1* | 2.365 | 3.300E-03 |
| *Lin7c* | 2.204 | 1.205E-03 |
| *Lmbrd2* | 2.736 | 4.057E-06 |
| *LOC310926* | -2.111 | 0.000E+00 |
| *LOC691921* | -2.456 | 7.946E-05 |
| *Lpgat1* | 2.282 | 6.139E-13 |
| *Lrrc23* | -4.418 | 2.195E-03 |
| *Ltbp2* | -2.575 | 2.598E-03 |
| *Map10* | 2.600 | 2.886E-02 |
| *Map2k4* | 2.283 | 3.208E-14 |
| *March1* | 3.118 | 1.286E-02 |
| *Mat2a* | 2.138 | 4.266E-09 |
| *Mbnl1* | 2.659 | 6.277E-05 |
| *Mctp2* | 4.270 | 1.244E-02 |
| *Mcu* | 2.627 | 8.220E-05 |
| *Med1* | 2.165 | 5.991E-05 |
| *Met* | 5.476 | 2.442E-04 |
| *Mfap3* | 2.016 | 3.602E-02 |
| *Mff* | 2.162 | 2.331E-02 |
| *Mfhas1* | 2.092 | 2.012E-02 |
| *Mfn2* | 3.586 | 4.587E-02 |
| *Mfsd4* | 2.557 | 1.926E-08 |
| *Mgat3* | 2.652 | 1.685E-15 |
| *Mgat5* | 2.062 | 2.966E-03 |
| *Mia* | -2.692 | 1.934E-02 |
| *Mid2* | 2.694 | 1.242E-02 |
| *Mir3561* | -2.132 | 4.611E-02 |
| *Mis12* | 2.370 | 2.817E-03 |
| *Mmp17* | 3.708 | 6.285E-14 |
| *Mn1* | 2.166 | 1.663E-03 |
| *Moap1* | 2.123 | 2.794E-03 |
| *Mob1b* | 2.858 | 5.158E-04 |
| *Mrgpre* | 2.151 | 2.357E-02 |
| *Mroh7* | -2.008 | 2.550E-04 |

| | | |
|---|---|---|
| *Msi1* | 2.100 | 3.731E-04 |
| *Mt1a* | -2.214 | 2.370E-06 |
| *Mtdh* | 2.376 | 2.199E-04 |
| *Mtmr9* | 2.223 | 2.396E-05 |
| *Mtpn* | 2.169 | 1.554E-24 |
| *Myt1l* | 2.200 | 3.614E-09 |
| *Napb* | 3.160 | 4.955E-16 |
| *Napepld* | 2.512 | 1.013E-03 |
| *Ndc1* | 2.313 | 4.085E-02 |
| *Necab1* | 2.098 | 3.370E-07 |
| *Negr1* | 3.096 | 1.856E-13 |
| *Neil2* | 3.310 | 4.534E-03 |
| *Nmnat2* | 2.008 | 3.668E-05 |
| *Npff* | -2.512 | 1.875E-02 |
| *Nrp1* | 2.022 | 1.471E-02 |
| *Nt5dc3* | 3.003 | 8.487E-05 |
| *Ntn4* | 2.142 | 1.513E-03 |
| *Nudcd3* | 3.169 | 2.370E-06 |
| *Nxph3* | 2.311 | 1.130E-06 |
| *Opa3* | 2.009 | 5.360E-10 |
| *Paics* | 2.170 | 3.370E-04 |
| *Pcdh11x* | 4.822 | 6.261E-03 |
| *Pcdh7* | 2.143 | 8.591E-08 |
| *Pcdha11* | 2.957 | 6.957E-03 |
| *Pcdha13* | 2.889 | 6.516E-03 |
| *Pcdhac2* | 2.423 | 1.078E-04 |
| *Pcdhga1* | 3.601 | 2.538E-03 |
| *Pcdhga10* | 2.842 | 2.889E-03 |
| *Pcdhga12* | 3.279 | 3.413E-05 |
| *Pcdhga2* | 4.238 | 1.179E-07 |
| *Pcdhga7* | 3.834 | 5.627E-07 |
| *Pcdhgb8* | 2.778 | 1.370E-02 |
| *Pcdhgc3* | 2.097 | 1.991E-03 |
| *Pclo* | 2.174 | 6.003E-12 |
| *Pcsk1* | 2.704 | 3.593E-02 |
| *Pcyox1* | 2.377 | 1.940E-18 |
| *Pde1a* | 2.604 | 3.575E-16 |
| *Pde4b* | 2.435 | 2.405E-13 |
| *Pdgfc* | 2.119 | 1.906E-02 |
| *Pdgfrl* | -2.183 | 1.508E-02 |
| *Pdlim5* | 2.147 | 1.263E-02 |
| *Pfkfb3* | 2.590 | 5.415E-05 |
| *Pfn2* | 2.213 | 2.176E-03 |
| *Pgam2* | 2.859 | 2.137E-02 |
| *Pgr* | 2.019 | 1.663E-03 |
| *Pi4k2a* | 2.339 | 1.650E-06 |
| *Pid1* | 2.311 | 4.022E-06 |
| *Pigb* | 2.846 | 1.745E-02 |
| *Pik3cb* | 2.526 | 2.211E-05 |

| | | |
|---|---|---|
| *Pip4k2b* | 2.201 | 1.405E-07 |
| *Pla2g2a* | -4.319 | 8.977E-03 |
| *Pnmal2* | 2.497 | 1.953E-32 |
| *Pofut1* | 2.220 | 1.488E-02 |
| *Polr2i* | -5.455 | 3.078E-03 |
| *Ppp2r2c* | 2.078 | 5.210E-07 |
| *Pramef8* | 2.231 | 7.291E-03 |
| *Prkaa2* | 3.081 | 6.875E-04 |
| *Prkacb* | 3.129 | 4.886E-20 |
| *Prkar2a* | 2.798 | 1.739E-04 |
| *Prkce* | 2.984 | 4.886E-20 |
| *Prrc2c* | 2.083 | 1.599E-05 |
| *Pter* | -2.078 | 5.280E-08 |
| *Rab11fip2* | 3.007 | 4.205E-05 |
| *Rab8b* | 2.006 | 3.769E-02 |
| *Rasa3* | 2.134 | 9.994E-06 |
| *Rasgrp1* | 3.044 | 1.964E-67 |
| *Rbfox3* | 2.266 | 1.094E-22 |
| *Rbm33* | 2.404 | 3.535E-04 |
| *Rc3h2* | 2.103 | 4.316E-02 |
| *Rell1* | 2.122 | 3.198E-02 |
| *Rfx3* | 2.606 | 1.164E-02 |
| *RGD1308601* | 3.033 | 2.952E-06 |
| *RGD1309821* | 2.410 | 9.548E-04 |
| *RGD1310262* | -2.542 | 4.218E-04 |
| *RGD1560436* | 3.111 | 2.221E-04 |
| *RGD1564124* | -2.092 | 2.165E-03 |
| *Rnf13* | 2.436 | 2.239E-04 |
| *Rnf24* | 2.525 | 6.635E-03 |
| *Rnf4* | 2.401 | 5.811E-09 |
| *Robo1* | 2.017 | 6.517E-03 |
| *Rpl3l* | -2.501 | 1.733E-02 |
| *Rps6* | -3.165 | 3.098E-04 |
| *Rrm2b* | 2.420 | 1.739E-02 |
| *Rrn3* | 3.117 | 5.678E-08 |
| *RT1-M1-2* | -2.057 | 2.853E-02 |
| *RT1-O1* | 2.863 | 1.426E-02 |
| *Rtkn2* | 3.128 | 1.365E-02 |
| *S100pbp* | 2.058 | 1.437E-04 |
| *Satb2* | 2.348 | 1.268E-09 |
| *Scamp5* | 2.651 | 4.359E-13 |
| *Scn2b* | 2.287 | 5.046E-09 |
| *Sel1l* | 2.102 | 5.400E-11 |
| *Senp2* | 2.117 | 6.755E-12 |
| *Sez6l* | 2.439 | 8.288E-21 |
| *Sh3gl2* | 2.111 | 6.472E-18 |
| *Slc14a1* | 2.451 | 6.270E-03 |
| *Slc16a14* | 2.435 | 1.914E-02 |
| *Slc24a2* | 2.840 | 5.670E-21 |

| | | |
|---|---|---|
| *Slc36a1* | 2.021 | 1.714E-02 |
| *Slc36a4* | 2.306 | 2.012E-05 |
| *Slc38a7* | 2.043 | 9.051E-04 |
| *Slc4a10* | 2.318 | 5.545E-26 |
| *Slc6a11* | 2.711 | 1.831E-05 |
| *Slc7a14* | 3.107 | 1.647E-04 |
| *Slco3a1* | 2.238 | 1.430E-05 |
| *Slk* | 2.334 | 1.438E-08 |
| *Slmap* | 2.234 | 7.699E-13 |
| *Smug1* | 3.417 | 2.048E-03 |
| *Sncg* | -3.826 | 9.470E-08 |
| *Snhg4* | -2.110 | 1.569E-04 |
| *Soat1* | 2.928 | 6.712E-05 |
| *Sobp* | 2.137 | 7.550E-07 |
| *Sphkap* | 2.314 | 1.909E-15 |
| *Spock2* | 2.570 | 1.165E-20 |
| *Spred2* | 2.158 | 5.134E-03 |
| *Sprn* | 2.664 | 1.931E-04 |
| *Spsb1* | 2.478 | 1.515E-04 |
| *Ss18l1* | 2.467 | 1.656E-03 |
| *St3gal1* | 2.689 | 1.189E-02 |
| *St6gal2* | 2.509 | 1.216E-02 |
| *St8sia4* | 2.414 | 1.508E-02 |
| *Stard7* | 2.148 | 3.777E-12 |
| *Stat4* | -2.193 | 1.228E-02 |
| *Stk26* | 4.030 | 2.856E-02 |
| *Stx1b* | 2.085 | 1.059E-15 |
| *Sulf1* | 2.023 | 3.164E-02 |
| *Syngr1* | 3.206 | 2.795E-14 |
| *Synpo* | 2.766 | 1.063E-22 |
| *Syt1* | 2.899 | 1.415E-28 |
| *Syt11* | 2.134 | 5.670E-21 |
| *Syt13* | 2.749 | 2.545E-18 |
| *Tacr1* | 2.639 | 4.289E-02 |
| *Tacr3* | 2.822 | 3.233E-10 |
| *Tarsl2* | 2.353 | 1.405E-07 |
| *Tbc1d24* | 2.156 | 7.764E-06 |
| *Tbr1* | 2.626 | 7.699E-13 |
| *Tbx19* | -3.710 | 9.056E-04 |
| *Tgfbr2* | 2.057 | 4.269E-03 |
| *Thap6* | 2.287 | 2.814E-03 |
| *Tmc8* | -2.376 | 3.332E-02 |
| *Tmem18* | 2.022 | 1.410E-04 |
| *Tmem196* | 2.297 | 4.082E-06 |
| *Tmem255a* | 3.508 | 1.374E-05 |
| *Tmtc2* | 2.602 | 1.023E-03 |
| *Tnik* | 2.030 | 1.585E-07 |
| *Tnmd* | -3.014 | 7.470E-03 |
| *Tomm70a* | 2.167 | 5.276E-09 |

| | | |
|---|---|---|
| *Trib1* | 2.738 | 1.007E-04 |
| *Tspyl4* | 2.265 | 6.408E-17 |
| *Ttc25* | -2.265 | 8.555E-03 |
| *Ttc4* | 2.270 | 1.038E-02 |
| *Tub* | 2.284 | 7.962E-04 |
| *Tubd1* | 2.398 | 2.342E-03 |
| *Ube2z* | 2.977 | 3.023E-11 |
| *Ubr1* | 2.099 | 5.949E-05 |
| *Unc5b* | 2.252 | 1.557E-03 |
| *Usp22* | 3.100 | 1.681E-20 |
| *Usp25* | 2.584 | 1.296E-07 |
| *Usp5* | 3.707 | 1.398E-07 |
| *Ust* | 2.664 | 1.251E-02 |
| *Vwc2l* | 2.329 | 9.515E-03 |
| *Wbp11* | 35.174 | 1.603E-02 |
| *Wbscr17* | 2.626 | 1.320E-08 |
| *Wdr54* | -6.249 | 2.263E-02 |
| *Wdr7* | 2.822 | 7.269E-14 |
| *Wipf2* | 2.151 | 1.791E-04 |
| *Wnt7b* | 3.171 | 2.332E-03 |
| *Wscd2* | 2.392 | 4.252E-07 |
| *Xk* | 4.757 | 3.574E-06 |
| *Xpr1* | 2.844 | 4.568E-05 |
| *Ypel2* | 3.004 | 1.904E-11 |
| *Zbtb38* | 2.297 | 7.855E-06 |
| *Zbtb46* | 2.349 | 3.215E-04 |
| *Zdhhc17* | 2.077 | 1.787E-11 |
| *Zfp238* | 2.266 | 1.347E-25 |
| *Zfp251* | 2.175 | 9.167E-06 |
| *Zfp319* | 2.662 | 9.009E-03 |
| *Zfp483* | 2.122 | 7.082E-03 |
| *Zfp516* | 3.481 | 7.057E-04 |
| *Zfp597* | 2.845 | 1.839E-04 |
| *Zkscan1* | 2.253 | 6.301E-04 |
| *Zmynd10* | -2.075 | 1.207E-03 |
