## Supplementary Material S.3 for "Repeated morphine exposure activates synaptogenesis and other neuroplasticity-related gene networks in the prefrontal cortex of male and female rats"

| Gene Symbol | Fold Change | FDR-adjusted p-value (q-value) |
| --- | --- | --- |
| *Acadsb* | 2.497 | 9.349E-06 |
| *Actr2* | 2.076 | 2.810E-06 |
| *Acvr1b* | 2.509 | 2.158E-07 |
| *Adam23* | 2.626 | 2.593E-05 |
| *Adamts2* | 2.090 | 1.050E-02 |
| *Aebp2* | 2.067 | 3.322E-03 |
| *Afap1* | 2.344 | 9.154E-03 |
| *Ahi1* | 2.066 | 2.777E-05 |
| *Akap5* | 2.202 | 2.154E-03 |
| *Alg10* | 2.498 | 2.510E-02 |
| *Amigo1* | 3.690 | 1.091E-07 |
| *Amph* | 2.691 | 4.189E-03 |
| *Ank1* | 2.017 | 2.984E-03 |
| *Ankrd34c* | 2.833 | 3.706E-04 |
| *Ankrd52* | 2.675 | 2.800E-03 |
| *Antxr1* | 2.008 | 1.298E-02 |
| *Ap2b1* | 2.038 | 2.235E-05 |
| *Ap4e1* | 2.662 | 6.454E-04 |
| *Apol9a* | -2.009 | 9.156E-03 |
| *Arhgap26* | 2.369 | 1.591E-02 |
| *Arhgap5* | 2.529 | 4.022E-06 |
| *Arid3a* | 2.714 | 2.922E-02 |
| *Arid5b* | 2.037 | 1.862E-02 |
| *Armc3* | -2.418 | 3.758E-02 |
| *Arrdc3* | 2.463 | 3.626E-03 |
| *Ate1* | 2.007 | 1.237E-02 |
| *Atf6* | 2.173 | 2.362E-03 |
| *Atp2b4* | 3.085 | 4.087E-05 |
| *Atp5f1* | 6.286 | 3.095E-02 |
| *Atxn3* | 2.048 | 2.382E-04 |
| *B3galt1* | 2.471 | 1.106E-03 |
| *Baz2a* | 2.031 | 1.394E-05 |
| *Bc1* | -2.247 | 1.055E-04 |
| *Bmp3* | 2.619 | 6.562E-03 |
| *Brpf3* | 2.300 | 9.773E-05 |
| *Bst2* | -2.455 | 4.316E-05 |
| *Btbd3* | 2.060 | 4.596E-04 |
| *C1galt1* | 2.630 | 7.964E-03 |
| *Cacna2d1* | 2.080 | 4.751E-04 |
| *Cacna2d2* | 2.429 | 5.816E-05 |
| *Cacnb4* | 2.090 | 1.449E-03 |
| *Cadps* | 2.168 | 2.404E-06 |
| *Caln1* | 2.051 | 1.993E-02 |
| *Camk1d* | 2.343 | 3.996E-06 |
| *Camk2g* | 4.342 | 7.326E-05 |
| *Capn5* | 2.190 | 3.426E-04 |
| *Carf* | 2.014 | 1.702E-03 |
| *Casc3* | 2.114 | 8.634E-05 |

| | | |
|---|---|---|
| *Cask* | 2.355 | 3.441E-02 |
| *Ccdc132* | 3.097 | 3.392E-08 |
| *Cd40* | -2.042 | 3.436E-02 |
| *Cdc14b* | 2.164 | 1.151E-02 |
| *Cdk18* | 2.084 | 1.294E-03 |
| *Cdk5r1* | 2.856 | 7.838E-05 |
| *Cds1* | 2.090 | 2.797E-04 |
| *Celsr2* | 2.263 | 5.790E-08 |
| *Cep85l* | 2.532 | 4.212E-02 |
| *Chid1* | -22.063 | 2.868E-03 |
| *Chml* | 3.344 | 3.280E-02 |
| *Chrna7* | 2.026 | 3.841E-02 |
| *Chrnb2* | 3.015 | 2.810E-06 |
| *Chst15* | 3.236 | 2.894E-03 |
| *Clcn2* | 28.741 | 3.647E-02 |
| *Clcn7* | 2.218 | 1.191E-04 |
| *Clint1* | 2.650 | 1.224E-05 |
| *Cnr2* | 2.840 | 2.830E-02 |
| *Coro2a* | 2.423 | 2.043E-02 |
| *Cox15* | 2.480 | 1.038E-03 |
| *Cpeb4* | 2.436 | 1.770E-05 |
| *Crim1* | 2.334 | 1.571E-05 |
| *Ctdspl2* | 2.373 | 2.155E-03 |
| *Ctif* | 3.022 | 4.301E-03 |
| *Cul5* | 2.068 | 1.407E-03 |
| *Cyth1* | 2.060 | 5.368E-03 |
| *Dcbld2* | 3.515 | 3.086E-06 |
| *Dck* | 2.175 | 4.162E-04 |
| *Dctn4* | 2.314 | 4.422E-05 |
| *Ddx6* | 2.452 | 2.810E-03 |
| *Dear* | 2.057 | 1.979E-04 |
| *Dgkg* | 2.789 | 6.436E-05 |
| *Dhcr24* | 2.630 | 7.917E-08 |
| *Diras2* | 2.337 | 5.016E-08 |
| *Dnajc16* | 2.406 | 1.149E-05 |
| *Drp2* | 2.099 | 1.226E-03 |
| *Dusp8* | 2.380 | 3.899E-05 |
| *Dvl3* | 2.077 | 4.913E-04 |
| *Edem3* | 2.602 | 2.161E-05 |
| *Ednrb* | 2.278 | 1.566E-03 |
| *Efcab2* | -2.341 | 1.989E-02 |
| *Efnb2* | 3.089 | 1.632E-05 |
| *Egr3* | 2.741 | 1.689E-02 |
| *Eogt* | 2.308 | 8.578E-03 |
| *Epha3* | 2.439 | 7.691E-04 |
| *Erc2* | 2.445 | 4.376E-05 |
| *Ern1* | 4.713 | 6.952E-03 |
| *Esrrg* | 2.276 | 2.703E-04 |
| *Etaa1* | 11.966 | 1.808E-05 |

| | | |
|---|---|---|
| *Etv3* | 2.286 | 3.334E-02 |
| *Exoc4* | 2.079 | 4.016E-04 |
| *Exoc8* | 2.120 | 5.411E-04 |
| *Extl3* | 2.525 | 3.374E-03 |
| *Eya3* | 2.193 | 1.803E-03 |
| *Fam134b* | 3.561 | 4.726E-03 |
| *Fam13a* | 2.245 | 3.885E-02 |
| *Fam168a* | 2.050 | 1.584E-07 |
| *Fam168b* | 2.242 | 2.252E-07 |
| *Fam217b* | 2.542 | 2.696E-04 |
| *Fam219a* | 2.011 | 1.101E-07 |
| *Fam49a* | 2.038 | 3.037E-03 |
| *Fam50a* | -3.090 | 6.189E-06 |
| *Fam65b* | 2.155 | 9.782E-05 |
| *Far1* | 2.073 | 3.123E-04 |
| *Fat4* | 2.527 | 3.031E-02 |
| *Fbxl16* | 2.009 | 1.947E-08 |
| *Ficd* | 2.061 | 2.249E-03 |
| *Fktn* | 2.078 | 4.082E-04 |
| *Flnb* | 3.534 | 5.891E-04 |
| *Fndc3b* | 2.782 | 4.320E-03 |
| *Fnip2* | 2.111 | 2.436E-02 |
| *Foxo1* | 2.358 | 4.422E-03 |
| *Foxo3* | 2.416 | 4.998E-04 |
| *Foxred2* | 4.601 | 5.601E-04 |
| *Foxs1* | -2.411 | 7.671E-04 |
| *Frmd4a* | 2.252 | 5.526E-03 |
| *Frmpd4* | 2.529 | 8.666E-06 |
| *Fry* | 2.252 | 1.900E-07 |
| *Fut10* | 3.941 | 1.692E-02 |
| *G3bp2* | 2.187 | 4.198E-05 |
| *Gabra4* | 2.388 | 1.214E-05 |
| *Gabrb2* | 2.815 | 3.037E-03 |
| *Gabrb3* | 2.313 | 2.857E-03 |
| *Galnt7* | 2.228 | 1.347E-02 |
| *Galntl6* | 2.184 | 1.288E-03 |
| *Gchfr* | -2.256 | 1.928E-02 |
| *Gda* | 2.893 | 2.662E-05 |
| *Glyr1* | 2.422 | 3.906E-06 |
| *Gmeb2* | 2.048 | 3.261E-02 |
| *Gna11* | 2.234 | 1.448E-04 |
| *Gnaq* | 2.281 | 2.070E-04 |
| *Gnl3l* | 2.338 | 9.926E-04 |
| *Gpd1l* | 2.295 | 1.225E-03 |
| *Gpr63* | 3.580 | 1.965E-02 |
| *Grid1* | 2.764 | 6.779E-04 |
| *Grin3a* | 2.466 | 4.886E-03 |
| *Grm5* | 2.736 | 3.111E-04 |
| *Hbb-b1* | -2.131 | 1.240E-04 |

| | | |
|---|---|---|
| *Hdac9* | 2.055 | 2.641E-02 |
| *Hiat1* | 3.577 | 4.179E-03 |
| *Hivep3* | 2.110 | 1.598E-03 |
| *Hsp90aa1* | 3.529 | 3.874E-03 |
| *Htr2a* | 2.608 | 7.084E-03 |
| *Htr5a* | 2.405 | 8.142E-03 |
| *Hunk* | 2.664 | 2.205E-03 |
| *Ifitm1* | -2.152 | 3.497E-04 |
| *Il10ra* | 2.421 | 1.612E-02 |
| *Il1rapl1* | 2.596 | 3.626E-02 |
| *Insr* | 2.041 | 3.070E-03 |
| *Ipcef1* | 2.529 | 1.749E-04 |
| *Irs1* | 2.312 | 6.521E-03 |
| *Isg15* | -2.865 | 1.831E-03 |
| *Itch* | 2.054 | 7.158E-05 |
| *Jph1* | 2.877 | 5.109E-05 |
| *Kalrn* | 2.319 | 2.709E-07 |
| *Kcnf1* | 2.292 | 3.229E-05 |
| *Kcnip2* | 3.377 | 4.082E-04 |
| *Kcnj13* | -2.125 | 7.698E-03 |
| *Kcnj3* | 2.292 | 9.755E-06 |
| *Kcnq2* | 2.644 | 4.548E-07 |
| *Klhl28* | 2.393 | 3.521E-03 |
| *Kpna5* | 2.841 | 8.885E-03 |
| *Kpnb1* | 2.101 | 3.231E-07 |
| *Ldlr* | 2.002 | 3.233E-04 |
| *Lhfpl4* | 3.049 | 6.282E-03 |
| *Lin7a* | 2.612 | 4.984E-02 |
| *Lin7c* | 2.264 | 3.096E-04 |
| *Lingo3* | 2.082 | 2.507E-02 |
| *Lman1l* | -2.410 | 3.238E-02 |
| *Lmbrd2* | 3.142 | 1.214E-05 |
| *LOC100910237* | -2.174 | 1.980E-02 |
| *LOC290595* | -2.590 | 2.045E-02 |
| *LOC310926* | -2.133 | 2.468E-06 |
| *LOC360231* | -2.737 | 9.581E-03 |
| *LOC367117* | -2.343 | 1.718E-02 |
| *LOC500034* | 2.313 | 2.459E-03 |
| *LOC501038* | 3.603 | 5.957E-03 |
| *Lpgat1* | 2.202 | 1.308E-04 |
| *Lrrc8b* | 2.282 | 4.452E-04 |
| *Lysmd3* | 2.291 | 4.193E-02 |
| *Lyst* | 2.061 | 1.626E-04 |
| *Magi1* | 2.057 | 1.158E-02 |
| *Mapk1ip1l* | 2.066 | 1.687E-02 |
| *Mapkbp1* | 2.748 | 1.882E-05 |
| *Mbnl1* | 2.238 | 9.079E-04 |
| *Mctp2* | 3.368 | 3.553E-03 |
| *Mcu* | 2.575 | 2.150E-04 |

| | | |
|---|---|---|
| *Med1* | 2.070 | 1.760E-04 |
| *Megf9* | 2.255 | 1.588E-05 |
| *Mei4* | -2.475 | 1.181E-02 |
| *Mfhas1* | 2.449 | 2.532E-03 |
| *Mfn2* | 3.295 | 1.388E-02 |
| *Mfsd4* | 2.510 | 6.129E-06 |
| *Mgat3* | 2.420 | 8.346E-06 |
| *Mgat5* | 2.265 | 7.795E-03 |
| *Mid2* | 2.643 | 1.167E-02 |
| *Mir3577* | 2.689 | 2.410E-02 |
| *Mmp17* | 3.442 | 2.911E-06 |
| *Mn1* | 2.025 | 3.261E-02 |
| *Moap1* | 2.155 | 6.041E-03 |
| *Mpp2* | 2.222 | 1.091E-07 |
| *Mrgpre* | 2.464 | 2.144E-03 |
| *Mroh1* | 2.723 | 1.226E-02 |
| *Mtdh* | 2.216 | 9.756E-04 |
| *Mtmr12* | 3.701 | 8.290E-05 |
| *Mtmr9* | 2.098 | 4.173E-03 |
| *Mx2* | -2.010 | 5.538E-07 |
| *Myo9a* | 2.129 | 1.999E-04 |
| *Myt1l* | 2.436 | 2.619E-05 |
| *Napb* | 3.286 | 2.148E-07 |
| *Nav3* | 2.110 | 2.810E-06 |
| *Ncoa2* | 2.148 | 5.694E-03 |
| *Ndc1* | 2.474 | 7.167E-03 |
| *Ndst3* | 2.031 | 1.322E-02 |
| *Necab1* | 2.348 | 2.797E-04 |
| *Negr1* | 2.544 | 2.404E-06 |
| *Nfix* | 2.160 | 6.436E-05 |
| *Nmb* | -2.293 | 1.634E-02 |
| *Nmnat2* | 2.107 | 2.617E-04 |
| *Noc3l* | 2.333 | 1.558E-04 |
| *Nos1ap* | 2.350 | 1.759E-07 |
| *Npff* | -2.834 | 8.162E-03 |
| *Nr4a3* | 2.174 | 8.564E-03 |
| *Nrp2* | 2.109 | 3.596E-02 |
| *Nt5dc3* | 3.274 | 3.382E-04 |
| *Nuak1* | 2.106 | 2.898E-03 |
| *Nudcd3* | 2.798 | 8.812E-04 |
| *Opcml* | 2.349 | 9.680E-04 |
| *Osbpl8* | 2.555 | 2.163E-05 |
| *Otx1* | 2.301 | 4.057E-02 |
| *Pacsin1* | 2.106 | 1.101E-06 |
| *Palm2* | 2.235 | 9.413E-03 |
| *Pamr1* | 2.116 | 1.970E-04 |
| *Pcdh11x* | 5.028 | 1.808E-05 |
| *Pcdhac2* | 3.108 | 3.486E-04 |
| *Pcdhga10* | 2.276 | 3.018E-03 |

| | | |
|---|---|---|
| *Pcdhga12* | 4.365 | 5.284E-05 |
| *Pcdhga2* | 2.270 | 3.322E-03 |
| *Pcdhga3* | 3.210 | 3.124E-02 |
| *Pcdhga5* | 3.226 | 1.198E-02 |
| *Pcdhga8* | 4.001 | 2.351E-04 |
| *Pcdhgb7* | 2.289 | 2.320E-03 |
| *Pcdhgb8* | 2.115 | 3.041E-02 |
| *Pcdhgc3* | 2.584 | 1.144E-05 |
| *Pdcd5* | -13.156 | 4.700E-06 |
| *Pde1a* | 2.303 | 3.852E-07 |
| *Pde4d* | 2.487 | 7.774E-03 |
| *Pdzd2* | 2.074 | 8.884E-03 |
| *Pfkfb3* | 2.542 | 2.021E-03 |
| *Pgr* | 2.337 | 9.036E-04 |
| *Phf21b* | 2.237 | 2.237E-02 |
| *Phlpp2* | 2.236 | 2.154E-03 |
| *Phyhip* | 18.102 | 2.116E-02 |
| *Pi4k2a* | 2.424 | 1.319E-04 |
| *Pik3cb* | 3.062 | 1.485E-05 |
| *Pnma2* | 2.082 | 3.906E-06 |
| *Pnmal2* | 2.546 | 1.759E-07 |
| *Pofut1* | 2.243 | 2.256E-02 |
| *Ppargc1a* | 2.246 | 6.115E-03 |
| *Ppm1e* | 2.148 | 3.195E-04 |
| *Ppp1r9a* | 2.106 | 1.621E-03 |
| *Ppp2r2c* | 2.031 | 7.937E-06 |
| *Prkaa2* | 2.222 | 3.511E-02 |
| *Prkacb* | 3.210 | 1.101E-06 |
| *Prkce* | 2.817 | 1.091E-07 |
| *Ptgs2* | 2.327 | 2.436E-03 |
| *Rab11fip2* | 3.440 | 6.075E-06 |
| *Rab8b* | 2.219 | 3.503E-02 |
| *Rad54l2* | 2.218 | 3.932E-03 |
| *Rapgef6* | 2.219 | 9.036E-04 |
| *Rasa3* | 2.186 | 4.557E-06 |
| *Rasgrp1* | 2.736 | 7.830E-11 |
| *Rbfox3* | 2.253 | 3.708E-07 |
| *Rbm33* | 2.177 | 2.984E-04 |
| *Reps1* | 2.206 | 2.070E-04 |
| *RGD1308601* | 2.254 | 7.216E-04 |
| *RGD1309821* | 2.227 | 1.143E-03 |
| *RGD1310429* | 2.056 | 1.186E-03 |
| *RGD1560289* | 2.018 | 3.931E-02 |
| *RGD1560436* | 2.461 | 4.522E-02 |
| *Rims2* | 2.035 | 3.328E-05 |
| *Rmrp* | -4.216 | 1.559E-03 |
| *Rnasel* | 2.296 | 1.071E-02 |
| *Rnf13* | 2.322 | 9.724E-06 |
| *Rnf144b* | 2.575 | 3.394E-04 |

| | | |
|---|---|---|
| *Rnf38* | 2.108 | 5.511E-03 |
| *Robo1* | 2.006 | 2.823E-03 |
| *Rpl17* | -2.635 | 0.000E+00 |
| *Rpl38* | -3.080 | 3.255E-04 |
| *Rps12* | -2.153 | 2.885E-02 |
| *Rps4y2* | -2.026 | 9.456E-03 |
| *Rrm2b* | 3.612 | 1.517E-04 |
| *Rrn3* | 2.710 | 4.618E-03 |
| *RT1-T24-3* | -2.044 | 2.893E-02 |
| *Rtkn2* | 3.627 | 1.831E-03 |
| *Rtp4* | -2.415 | 2.077E-03 |
| *S100a11* | -2.284 | 5.050E-03 |
| *S100a6* | -2.101 | 1.186E-03 |
| *S100pbp* | 2.083 | 2.284E-04 |
| *Sall3* | 2.193 | 6.226E-03 |
| *Satb2* | 2.364 | 1.616E-04 |
| *Scamp5* | 2.610 | 1.101E-06 |
| *Scn1a* | 2.192 | 1.453E-03 |
| *Scn2b* | 2.698 | 2.070E-04 |
| *Scn8a* | 2.107 | 2.070E-04 |
| *Scrn3* | 2.281 | 1.110E-03 |
| *Sel1l* | 2.135 | 1.015E-05 |
| *Sez6l* | 2.457 | 2.023E-05 |
| *Sgms2* | -2.212 | 3.982E-02 |
| *Sgpp2* | 2.663 | 1.932E-03 |
| *Sh3pxd2a* | 2.207 | 7.473E-04 |
| *Sik1* | 2.124 | 2.726E-02 |
| *Ska1* | -2.654 | 4.572E-02 |
| *Slc14a1* | 2.704 | 1.373E-03 |
| *Slc16a14* | 2.012 | 1.825E-02 |
| *Slc1a4* | 2.282 | 7.496E-04 |
| *Slc24a2* | 2.799 | 7.830E-11 |
| *Slc25a15* | 2.439 | 4.130E-03 |
| *Slc35e2b* | 2.218 | 3.088E-02 |
| *Slc36a1* | 2.967 | 1.699E-03 |
| *Slc36a4* | 2.338 | 4.537E-04 |
| *Slc39a11* | 2.117 | 1.069E-02 |
| *Slc39a9* | 2.093 | 1.591E-02 |
| *Slc4a7* | 2.996 | 4.747E-03 |
| *Slc6a11* | 2.136 | 1.449E-03 |
| *Slc6a5* | 2.195 | 2.585E-02 |
| *Slc6a6* | 2.396 | 1.706E-04 |
| *Slc9a8* | 2.669 | 2.606E-04 |
| *Slfn3* | -2.782 | 5.467E-03 |
| *Slk* | 2.284 | 1.250E-04 |
| *Slmap* | 2.090 | 1.476E-05 |
| *Smo* | 2.010 | 5.306E-03 |
| *Smug1* | 3.313 | 2.128E-03 |
| *Snrk* | 2.182 | 3.169E-04 |

| | | |
|---|---|---|
| *Sp4* | 2.674 | 8.290E-05 |
| *Sphkap* | 2.048 | 4.528E-04 |
| *Spock2* | 2.625 | 5.053E-06 |
| *Spred2* | 2.399 | 1.302E-04 |
| *Spsb1* | 3.008 | 1.038E-05 |
| *Sptb* | 3.157 | 3.830E-06 |
| *Ss18l1* | 2.574 | 8.687E-04 |
| *Ssh2* | 2.005 | 6.553E-04 |
| *Strn* | 2.166 | 1.142E-03 |
| *Stx1b* | 2.349 | 3.852E-07 |
| *Syngr1* | 3.111 | 2.158E-07 |
| *Synpo* | 2.472 | 1.144E-05 |
| *Syt1* | 2.637 | 5.999E-08 |
| *Syt11* | 2.140 | 2.158E-07 |
| *Syt13* | 2.434 | 1.190E-06 |
| *Syt7* | 2.077 | 1.319E-04 |
| *Tacr1* | 2.748 | 3.625E-02 |
| *Tacr3* | 2.207 | 1.803E-04 |
| *Tanc2* | 2.169 | 7.427E-05 |
| *Tap2* | -2.146 | 3.991E-02 |
| *Tbc1d24* | 2.422 | 4.314E-05 |
| *Tbc1d7* | 2.039 | 1.644E-02 |
| *Tbcel* | 2.241 | 2.800E-03 |
| *Tbr1* | 2.178 | 7.838E-05 |
| *Tgfbr3* | 2.147 | 4.808E-03 |
| *Thap6* | 2.284 | 8.735E-03 |
| *Timp1* | -2.022 | 1.069E-02 |
| *Tmem132d* | 2.223 | 1.558E-04 |
| *Tmem196* | 2.100 | 4.712E-04 |
| *Tmem255a* | 4.225 | 1.858E-04 |
| *Tmem87b* | 2.018 | 3.968E-02 |
| *Tmtc2* | 2.183 | 3.424E-03 |
| *Tom1l2* | 2.143 | 9.856E-05 |
| *Tomm70a* | 2.225 | 2.911E-06 |
| *Trib1* | 2.911 | 1.659E-02 |
| *Tsen34* | 188.815 | 2.076E-02 |
| *Tspyl4* | 2.209 | 1.144E-05 |
| *Ttc4* | 2.582 | 4.881E-04 |
| *Tub* | 2.430 | 7.878E-04 |
| *Twsg1* | 3.267 | 1.448E-04 |
| *Ube2z* | 2.104 | 5.557E-04 |
| *Usp22* | 2.228 | 1.006E-03 |
| *Usp25* | 2.436 | 4.257E-04 |
| *Usp5* | 3.118 | 1.144E-05 |
| *Uvrag* | 2.215 | 1.226E-03 |
| *Vwc2l* | 2.159 | 3.461E-02 |
| *Wbp11* | 37.025 | 2.205E-02 |
| *Wbscr17* | 2.482 | 7.216E-04 |
| *Wdfy1* | 2.239 | 7.088E-04 |

| | | |
|---|---|---|
| *Wdr7* | 2.946 | 5.992E-06 |
| *Wipf2* | 2.452 | 9.670E-05 |
| *Wscd2* | 2.907 | 6.189E-06 |
| *Ypel2* | 2.405 | 8.233E-05 |
| *Zbtb16* | 2.287 | 1.928E-02 |
| *Zbtb25* | 2.559 | 1.073E-03 |
| *Zbtb26* | 2.191 | 3.314E-02 |
| *Zbtb38* | 2.037 | 6.557E-03 |
| *Zdhhc17* | 2.122 | 1.091E-07 |
| *Zdhhc23* | 2.924 | 1.295E-04 |
| *Zfp251* | 2.047 | 1.831E-03 |
| *Zfp319* | 2.820 | 6.454E-04 |
| *Zfp382* | 2.035 | 1.166E-04 |
| *Zfp483* | 2.750 | 4.848E-05 |
| *Zfp516* | 3.177 | 1.441E-02 |
| *Zfp597* | 3.187 | 1.750E-03 |
| *Zfp707* | 2.287 | 1.820E-02 |
| *Zkscan1* | 2.097 | 2.477E-03 |
