## Supplementary Material S.4 for "Repeated morphine exposure activates synaptogenesis and other neuroplasticity-related gene networks in the prefrontal cortex of male and female rats"

| Gene Symbol | Assay ID | Gene Name | RNA-seq | | | | RT-qPCT | | | |
|---|---|---|---|---|---|---|---|---|---|---|
| | | | Male | | Female | | Male | | Female | |
| | | | Fold-change | FDR adj. *p*-value | Fold-change | FDR adj. *p*-value | Fold-change | *p*-value | Fold-change | *p*-value |
| *Cdk5r1* | Rn02132948_s1 | cyclin-dependent kinase 5, regulatory subunit 1 (p35) | 2.475 | 0.0000001 | 2.856 | 0.0000784 | 1.461 | 0.01327* | 0.992 | 0.9403 |
| *Chrnb2* | Rn00570733_m1 | cholinergic receptor, nicotinic, beta 2 (neuronal) | 2.554 | 0.0000000 | 3.015 | 0.0000028 | 1.182 | 0.1114 | 1.045 | 0.6815 |
| *Efnb2* | Rn01756899_m1 | ephrin B2 | 6.279 | 0.0000000 | 3.089 | 0.0000163 | 1.157 | 0.1053 | 1.080 | 0.3725 |
| *Esrrg* | Rn01415309_m1 | estrogen-related receptor gamma | 2.201 | 0.0009110 | 2.276 | 0.0002700 | 1.083 | 0.1006 | 1.153 | 0.2569 |
| *Gabra4* | Rn00589846_m1 | gamma-aminobutyric acid (GABA) A receptor, alpha 4 | 2.551 | 0.0000000 | 2.388 | 0.0000121 | 1.077 | 0.2087 | 1.098 | 0.2641 |
| *Gabrb2* | Rn00564149_m1 | gamma-aminobutyric acid (GABA) A receptor, beta 2 | 2.286 | 0.0024419 | 2.815 | 0.0030370 | 1.187 | 0.03267* | 1.007 | 0.9391 |
| *Grid1* | Rn01453420_m1 | glutamate receptor, ionotropic, delta 1 | 2.658 | 0.0000001 | 2.764 | 0.0006780 | 1.043 | 0.5613 | 1.199 | 0.03497* |
| *Grm5* | Rn00566628_m1 | glutamate receptor, metabotropic 5 | 2.326 | 0.0000005 | 2.736 | 0.0003110 | 1.244 | 0.02307* | 1.071 | 0.3752 |
| *Htr5a* | Rn00565746_m1 | 5-hydroxytryptamine (serotonin) receptor 5A, G protein-coupled | 2.200 | 0.0009095 | 2.405 | 0.0081421 | 1.057 | 0.5017 | 1.089 | 0.4981 |
| *Negr1* | Rn00572380_m1 | neuronal growth regulator 1 | 3.096 | 0.0000000 | 2.544 | 0.0000024 | 1.131 | 0.3011 | 1.195 | 0.0830 |
| *Pgr* | Rn01448227_m1 | progesterone receptor | 2.019 | 0.0016600 | 2.337 | 0.0009040 | 1.127 | 0.0608 | 1.167 | 0.0428* |
| *Rtp4* | Rn01404500_m1 | receptor (chemosensory) transporter protein 4 | - | - | -2.415 | 0.0020800 | - | - | 0.897 | 0.7192 |
| *Sncg* | Rn00581652_m1 | synuclein, gamma | -3.826 | 0.0000001 | - | - | 0.674 | 0.3765 | - | - |
| *Actb* | Rn00667869_m1 | actin, beta | Endogenous control | | | | | | | |
