## Supplementary Material S.5 for "Repeated morphine exposure activates synaptogenesis and other neuroplasticity-related gene networks in the prefrontal cortex of male and female rats"

| Upstream Regulator | Molecule Type | Predicted Activation State | Activation z-score | p-value of overlap |
|---|---|---|---|---|
| JAK1/2 | group | Activated | 2 | 0.00307 |
| FEV | transcription regulator | Activated | 2 | 0.00666 |
| ADORA2A | G-protein coupled receptor | Activated | 2 | 0.0267 |
