## Supplementary Material S.6 for "Repeated morphine exposure activates synaptogenesis and other neuroplasticity-related gene networks in the prefrontal cortex of male and female rats"

| Ingenuity Canonical Pathways | -log(p-value) | z-score |
|---|---|---|
| Synaptogenesis Signaling Pathway | 8.23 | 4.264 |
| Neuroinflammation Signaling Pathway | 4.59 | 2.53 |
| cAMP-mediated signaling | 4.16 | 3.051 |
| AMPK Signaling | 3.82 | 2.121 |
| Synaptic Long Term Potentiation | 2.99 | 2.828 |
| Calcium Signaling | 2.78 | 3 |
| Opioid Signaling Pathway | 2.67 | 3.317 |
| Ephrin Receptor Signaling | 2.64 | 2.236 |
| Gαq Signaling | 2.44 | 2.828 |
| Dopamine-DARPP32 Feedback in cAMP Signaling | 2.36 | 2.449 |
| CREB Signaling in Neurons | 1.75 | 2.646 |
| Ephrin B Signaling | 1.57 | 2 |
| Gαs Signaling | 1.57 | 2.236 |
| G Beta Gamma Signaling | 1.36 | 2.236 |
