## Supplementary Material S.7 for "Repeated morphine exposure activates synaptogenesis and other neuroplasticity-related gene networks in the prefrontal cortex of male and female rats"

| Ingenuity Canonical Pathways | -log(p-value) | z-score |
|---|---|---|
| Synaptogenesis Signaling Pathway | 6.25 | 4.243 |
| Endocannabinoid Neuronal Synapse Pathway | 4.87 | 2.714 |
| AMPK Signaling | 4.68 | 2.121 |
| Calcium Signaling | 4.21 | 2.714 |
| G Beta Gamma Signaling | 2.9 | 2.828 |
| CREB Signaling in Neurons | 2.48 | 2.236 |
| Opioid Signaling Pathway | 2.38 | 3.317 |
| Synaptic Long Term Potentiation | 2.15 | 2.646 |
| Synaptic Long Term Depression | 1.8 | 2.121 |
| cAMP-mediated signaling | 1.77 | 2.828 |
| Ephrin B Signaling | 1.45 | 2 |
| Ephrin Receptor Signaling | 1.44 | 2.236 |
