## Supplementary Material Captions for "Repeated morphine exposure activates synaptogenesis and other neuroplasticity-related gene networks in the prefrontal cortex of male and female rats"

S.1 Principal Component Analysis of the 500 most divergent genes. Labels indicate samples used for RNA-seq: Blue, Male/Morphine; Orange, Male/Saline; Green, Female/Morphine; Red, Female/Saline. Dimension 1, drug treatment; Dimension 2, sex.

S.2 Male differentially expressed gene list [Absolute Fold Change > 2, false discovery rate (FDR) corrected $p$-value ($q$-value) $< 0.05$, and > 20 reads for at least one of the experimental conditions].

S.3 Female differentially expressed gene list [Absolute Fold Change > 2, false discovery rate (FDR) corrected $p$-value ($q$-value) $< 0.05$, and > 20 reads for at least one of the experimental conditions].

S.4 RT-qPCR probe ID and full results (n = 4-6/group).

S.5 IPA analysis of upstream regulators derived from the set of differentially expressed genes common to males and females (absolute $z$-score $\geq 2$; $p$-value $< 0.05$).

S.6 IPA analysis of affected canonical pathways with male dataset [absolute $z$-score $\geq 2$; $p$-value $< 0.05$, $-\log(p\text{-value}) > 1.3$].

S.7 IPA analysis of affected canonical pathways with female dataset [absolute $z$-score $\geq 2$; $p$-value $< 0.05$, $-\log(p\text{-value}) > 1.3$].
